# Dissecting the allosteric networks governing agonist efficacy and potency in G protein-coupled receptors

**DOI:** 10.1101/2021.09.14.460253

**Authors:** Franziska M. Heydenreich, Maria Marti-Solano, Manbir Sandhu, Brian K. Kobilka, Michel Bouvier, M. Madan Babu

**Author notes:** These authors contributed equally (Maria Marti-Solano, Manbir Sandhu).

## Abstract

G protein-coupled receptors (GPCRs) translate binding of extracellular ligands into intracellular responses through conformational changes. Ligand properties are described by the maximum response (efficacy) and the agonist concentration at half-maximal response (potency). Integrating structural changes with pharmacological properties remains challenging and has not yet been performed at the resolution of individual amino acids. We use epinephrine and β2-adrenergic receptor as a model to integrate residue-level pharmacology data with intramolecular residue contact data describing receptor activation. This unveils the allosteric networks driving ligand efficacy and potency. We provide detailed insights into how structural rearrangements are linked to fundamental pharmacological properties at single-residue level in a receptor-ligand system. Our approach can be used to determine such pharmacological networks for any receptor-ligand complex.

## Introduction

G protein-coupled receptors (GPCRs) form a major family of membrane proteins that respond to a variety of extracellular ligands including photons, small molecules neurotransmitters, and hormones (*1-5*). In humans, over 500 endogenous ligands (*6*) act on ∼350 non-odorant GPCRs and regulate many aspects of human physiology (*7, 8*). Given their capacity to modulate human physiology, about one-third of all FDA approved drugs target one of these GPCRs to treat various diseases (*9*). During receptor signaling, ligand binding is converted into an intracellular signal via conformational changes in the GPCR (*10*). The ligand shifts the conformational equilibrium of the receptor towards active or inactive states depending on whether it is an agonist or an inverse agonist (*11*). The conformational states associated with agonist binding strongly increase the likelihood of signaling response (e.g., G protein activation). Signaling response to a ligand that targets a receptor is characterized by efficacy and potency (*12*); two fundamental pharmacological properties that are measured by monitoring the signaling response of the receptor to varying ligand concentrations (i.e., dose/concentration-response curves). Efficacy relates to the maximum amplitude of a signaling response, whereas potency refers to the agonist concentration where signaling reaches the half-maximal response. Given their importance in describing such systems, efficacy and potency have been characterized for numerous ligand-receptor systems for several decades (*13*). In the case of GPCRs and their endogenous agonists, the amino acid sequence of a receptor and its 3D structure have been subject to strong selection pressure so that ligand efficacy and potency are within physiologically tolerable range. Synthetic ligands that are used as drugs possess their own combination of efficacy and potency values that is determined by the way in which they specifically engage with receptor structure.

Despite thorough characterization of numerous ligand-GPCR pairs over the years, we do not yet fully understand the molecular determinants of efficacy and potency for any agonist-GPCR pair. Little is known about how agonist efficacy and potency are determined by receptor sequence and structure at the resolution of single amino acids. For a given agonist-receptor pair, which residues in the receptor determine efficacy and potency for a signaling response? What is the spatial relationship between those residues in an active conformational state and with respect to key functional regions such as agonist and effector binding sites? Addressing these questions would allow us to discriminate auxiliary structural changes from those that are functionally important. It would also enable us to determine the critical residues for signal transmission (i.e., drivers) and the underlying allosteric networks associated with efficacy and potency. To identify how efficacy and potency are encoded in receptor sequence and structure, we designed a study to infer the main receptor determinants governing these properties. We used the β2-adrenergic receptor (β2AR), a well-studied, prototypical Family A GPCR involved in smooth muscle relaxation. In humans, β2AR is known to be activated by epinephrine (adrenaline) and to signal through the heterotrimeric G protein, Gs. By developing a data science framework that integrates (i) experimentally measured values of epinephrine efficacy and potency towards Gs signaling upon mutation of every position in β2AR and (ii) data on structural changes associated with agonist binding, receptor activation and Gs coupling (**Fig. 1**), we reveal how ligand efficacy and potency are encoded in receptor sequence and structure in a prototypical GPCR.

**Fig. 1.**
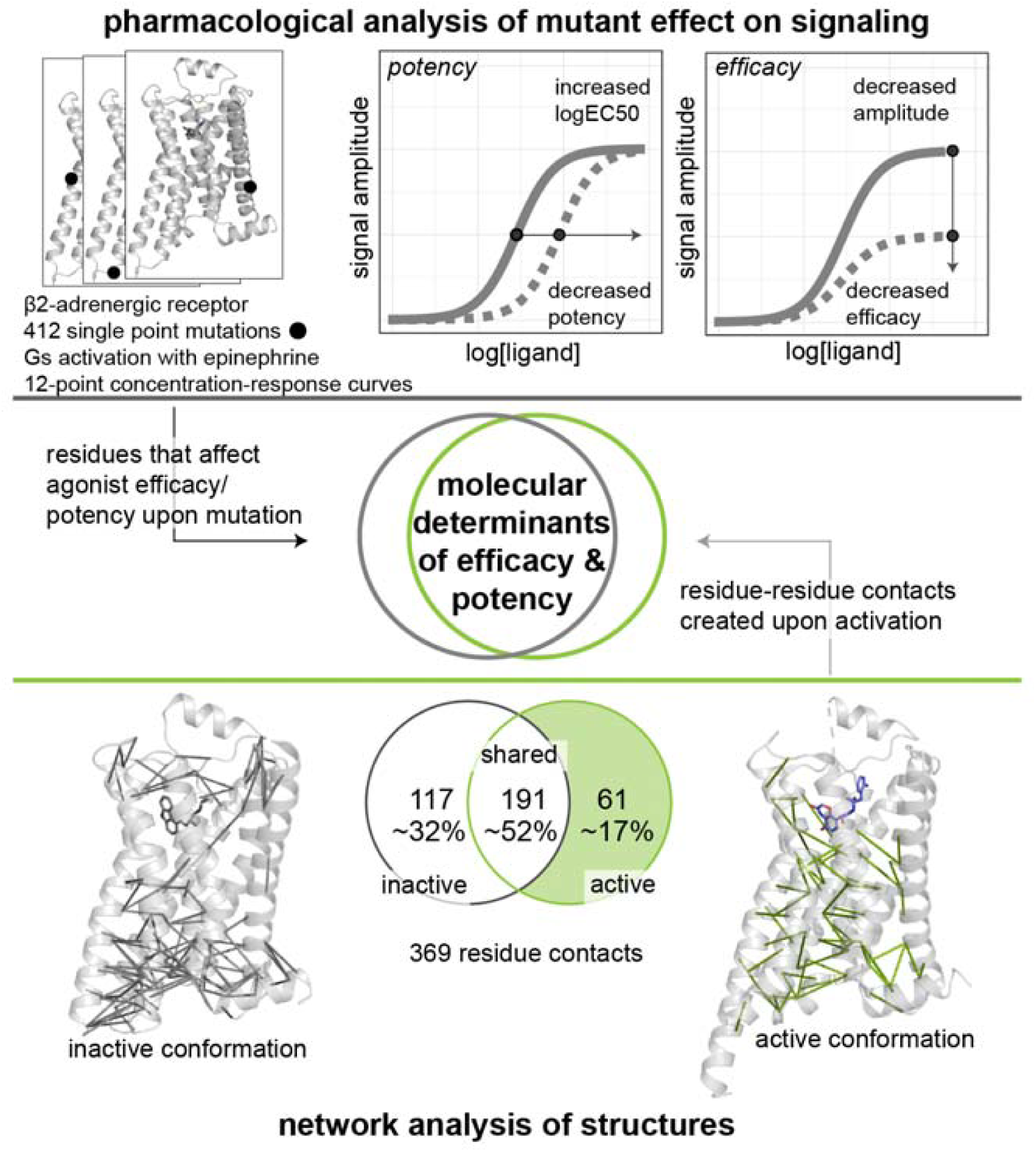
An approach to integrate pharmacological and structural data to reveal the molecular determinants and allosteric networks of efficacy and potency. Every residue of a receptor is mutated and key pharmacological properties for a ligand-receptor-system is determined (top panel). Active and inactive states of receptors are analyzed as networks of non-covalent contacts to infer those that are active state specific (bottom panel). These data are integrated to discover the molecular determinants and the underlying allosteric networks of efficacy and potency (middle panel).

## Results

### A restricted number of β2AR residues drive epinephrine efficacy and potency

First, we investigated which positions in β2AR are important for agonist efficacy and potency. We created 412 single-point mutants of β2AR, one for each position in the receptor, and evaluated their Gs signaling profile by obtaining concentration-response curves using its endogenous agonist, epinephrine. To minimize secondary structure disturbances to the secondary structure of the receptor, we replaced each native amino acid with alanine, or glycine if the native amino acid was alanine. Both types of substitutions are well tolerated in the long, membrane-spanning α-helices (*14*) of a GPCR. To facilitate comparison across receptors, we refer to residues using the GPCRdb numbering scheme (e.g. L124 as ‘3×43’) (*15*), a system based on Ballesteros-Weinstein numbering (*16*) wherein the first number refers to the helix or loop and the second number refers to the position relative to the most conserved residue in that helix.

We measured the effect of each β2AR mutation on the activation of heterotrimeric Gs in response to epinephrine in a cell-based assay using a BRET (Bioluminescence resonance energy transfer)-based biosensor (*17, 18*) (**Fig. S1**, see **Methods)**. Briefly, the biosensor reports on the distance between the α and γ subunits of the G protein; this distance increases upon receptor activation due to conformational changes in the α subunit and its dissociation from the βγ-subunits. Each assay was performed as 12-point concentration-response curves in biological triplicates (**Fig. 1, top panel, Methods**) to determine the two parameters in the Gs signaling response: the signal amplitude (efficacy) and logEC50 (potency) (**Fig. S1**). We also performed measurements on cells expressing wild type β2AR or no receptor as controls. In total, we obtained over 16,000 measurements and the parameters for agonist induced Gs activation are reported in **Table S1**.

Since low cell-surface expression negatively influences the measurement of signaling, we also evaluated the expression levels of all mutants by cell-surface ELISA (**Fig. S2A-C**, see **Methods**). For expression levels under 25% of wild type, one cannot discriminate whether the observed reduction in signaling is due to the altered function of the mutant or due to reduced receptor expression (**Fig. S2A-B**). Thus, the signal measurements of mutants in this expression regime were deemed unreliable. We thus excluded 16 mutations with low expression (<25% of WT level) from the pharmacological analysis (**Fig. S2C**). Of those 16 mutations, two mapped to the first transmembrane (TM) helix (N51^1×50^ and I55^1×54^), suggesting that they could be critical for membrane protein folding and biogenesis in the endoplasmic reticulum (ER) membrane. One mutation mapped to the intracellular loop 1 (ICL1; L64^12×50^), a residue previously identified as important for ER export (*19*). Six residues made two distinct sets of contacts with each other in the structure. The first set involved a conserved disulfide bridge and a neighboring tryptophan, recently shown to be particularly intolerant to mutation (*20*) (cf. **Fig. S2D-E**). The second set of contacts involved residues in TM 2, 6 and 7 via the highly conserved W^6×48^, part of the CWxP motif consisting of residues C/S/T^6×47^, W^6×48^ and P^6×50^ at the bottom of the ligand binding pocket; their distribution across the TM helices suggests their likely importance to maintain the fold of the helix bundle in the membrane.

The other 396 mutants showed cell-surface expression with values >25% of the wild-type β2AR and were therefore considered further for the pharmacological analysis. We discretized the efficacy and potency values (i.e., classified as affected/not affected) to discover patterns that can help explain the pharmacological response through data science approaches. We defined cut-offs to distinguish positions where mutations showed a wild-type-like signaling profile from positions where mutations strongly affected signaling (see **Methods**). Cut-offs were chosen to control for the effect of the expression level and signaling response. i.e., decreased signaling is not explained by decreased expression level. We found that most (>75%) single-point mutations did not affect Gs signaling. 82 mutations satisfied the cut-off and were therefore classified as strongly reducing receptor signaling. Of these 82 mutations, 3 led to no measurable Gs activation, 21 only reduced efficacy, 37 only reduced potency, while 21 reduced both potency and efficacy (**Fig. 2A**). These findings suggest that efficacy and potency of the endogenous agonist epinephrine are likely governed by a subset of residues (∼20%), which are thus pharmacological determinants for this ligand.

**Fig. 2.**
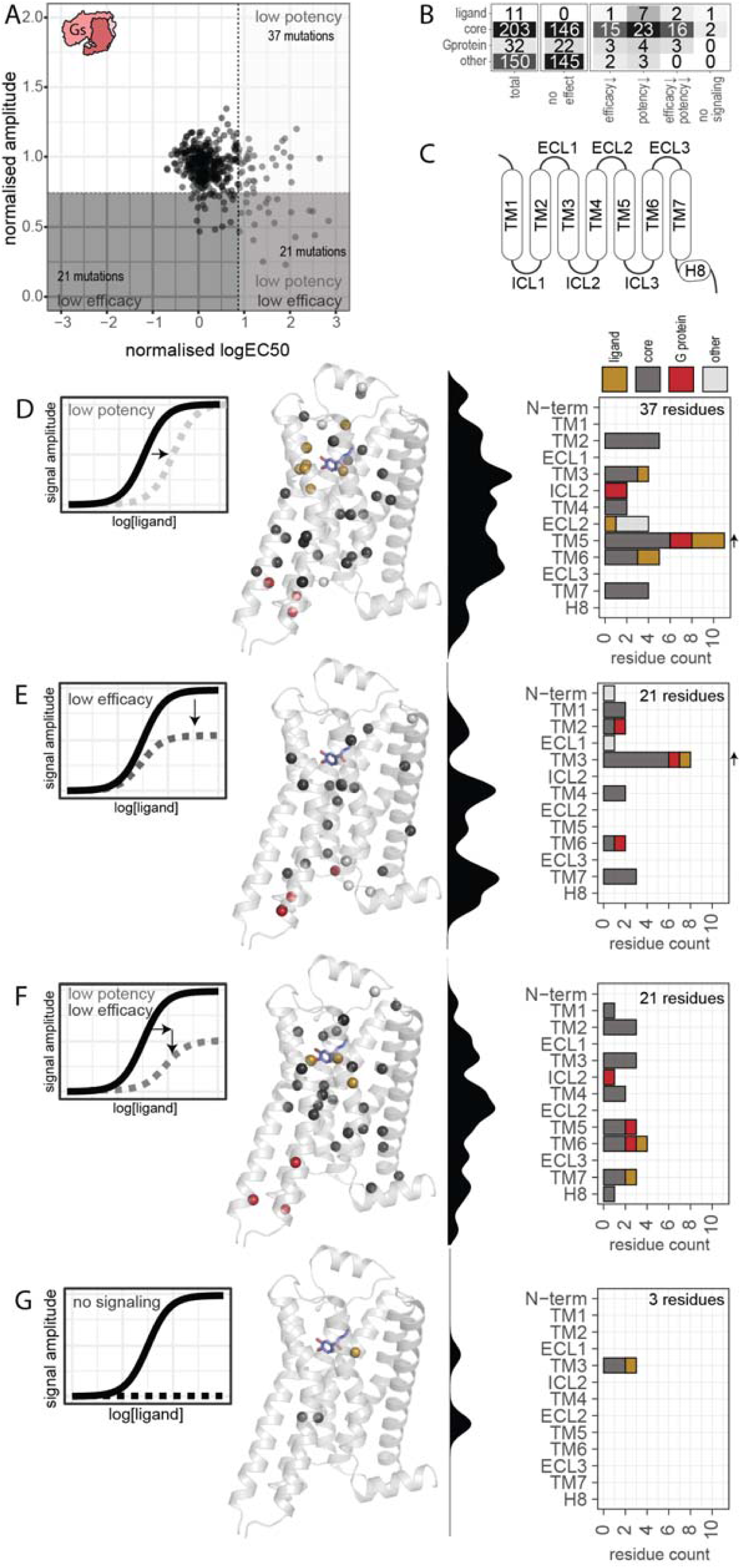
Receptor positions that strongly affect efficacy or potency upon mutation. (**A**) Overview of potency (normalized logEC50) and efficacy (normalized signal amplitude) of all β2AR mutations. Cut offs are shown as dashed lines. Mutations that affected potency and/or efficacy compared to the wild type are marked with grey boxes. (**B**) Distribution of residues by functional regions and their effect on efficacy and potency. Ligand-binding pocket and G protein–binding sites include residues within 4 Å of epinephrine (PBD 4LDO) or Gs (PDB 3SN6). (**C**) A cartoon representation of β2AR showing the position of secondary structure elements. Transmembrane helices are numbered. (**D–G**) Signaling profile, position of mutated residue on the structure (PDB 4LDO), the density of mutations on the extra-to intracellular axis and a bar graph of the location of mutation in relation to its secondary structure element and function regions is shown for mutations with decreased potency (D), decreased efficacy (E), decreased potency and efficacy (F), and mutations that decreased signaling to a non-measurable level (G). Arrows indicate statistical overrepresentation of mutations with a certain effect in the secondary structure element (**Methods**).

We examined where these positions lie on the receptor structure with respect to the various secondary structure elements (**Fig. 2B-G**). In other words, are they distributed throughout the structure, or do individual secondary structure elements act as key determinants of efficacy and potency? To address this question, we developed a computational approach to combine the pharmacological information on how each mutation affected the efficacy and potency of epinephrine for Gs signaling with sequence and 3D structure data of the receptor (**Fig. 1, middle and bottom panel; Methods**). We observed that TM5 was enriched in residues that affected potency only, whereas TM3 was enriched in residues that affect efficacy only. In contrast, the positions that affected efficacy and potency simultaneously were distributed across the different transmembrane helices. This observation suggests that distinct secondary structure elements contribute to efficacy and potency, while residues that affect both properties are not localized to any single secondary structure element.

We then investigated the relationship of the residues that affect efficacy and potency with respect to the key functional regions such as the ligand-binding pocket, the receptor core, and the G protein-binding cavity (**Fig. 2B**). In order to do so, we first analyzed existing structures of β2AR to identify 32 residues that contact the G protein (PDB 3SN6 and 6E67) and 11 residues that contact epinephrine (PDB 4LDO, (*21*)) (**Methods**); all other residues in the transmembrane helices were defined as core residues (203 residues). Only 10 of the 32 G protein-contacting residues, showed an effect when mutated. 4 positions only affected potency (P138^34×50^, Y141^34×53^, I233^5×72^, V222^5×61^), 3 only affected efficacy (T68^2×39^, I135^3×54^, E268^6×30^) and 3 affected both (F139^34×51^, A226^5×65^, L275^6×37^; **Fig. S3**). These results suggest that (a) a large fraction of residues contacting the G protein are evolvable (i.e., can mutate) with little effect on efficacy or potency and (b) there are hot spots in the G protein interface that have a distinct effect on efficacy and potency. In contrast, all 11 epinephrine-contacting residues negatively affected efficacy and/or potency when mutated (**Fig. S4**). Most (7 of 11) positions affected only potency when mutated and one mutation completely abolished signaling. The latter residue, D113^3×32^A, is known to be critical for agonist binding in β2AR (*22, 23*) and other aminergic receptors, as it contacts the positively charged amine group that is a common feature of endogenous agonists for monoamine receptors.

### Residues driving efficacy and potency form allosteric contact networks

The pattern of distribution of the pharmacological determinants of receptor signaling raises several questions about the spatial relationship between the residues that affect efficacy and/or potency. Do they form allosteric networks of residue contacts, and if so, do the residue contact networks that govern efficacy and potency spatially overlap or segregate? While it is well appreciated that the receptor undergoes major structural changes upon activation (*24-27*), we do not understand which of the rearrangements, and which residues that change conformation are directly relevant for the efficacy and potency of the receptor response. To address these questions, we analyzed active and inactive state structures to define residue contact networks. In such networks, each residue is represented as a node and non-covalent contacts between residues are represented as edges or links. We compared structures of β2AR in inactive and active states to identify residues that participate in active state–specific contacts (**Fig. S5**). For the inactive state, we chose the highest resolution structure available (with antagonist carazolol, 2.4 Å, PDB 2RH1), whereas for the active state we analyzed the only available structure of β2AR in complex with Gs (with the agonist BI-167107, 3.2 Å, 3SN6)(*27-29*).

We then integrated the experimentally determined pharmacological properties with the residue contact network (**Fig. 1, middle panel**; **Methods**). Integration of these orthogonal data provides a new perspective allowing us to stratify residues based on their importance for pharmacology and for structural rearrangement upon receptor activation (**Fig. 3A**). We first classified every mutant as *affected* (i.e., if it affects efficacy, potency, or both) or *non-affected* (no effect on efficacy, potency, or both). We then identified all residue contacts that are present only in the active state. Based on whether they affected pharmacology and/or mediate an active state specific contact or not, we defined: (a) “driver residues” mediate an active-state specific contact with another residue that similarly affect pharmacology when mutated, (b) “modulator residues” are non-driver residues that affect pharmacology when mutated; (c) “passenger residues” mediate an active-state specific contact but have no effect on pharmacology when mutated and (d) “bystander residues” do not mediate active-state specific contacts and have no effect on pharmacology when mutated (**Methods)**. This classification allowed us, for the first time, to describe how potency and efficacy signals are allosterically transmitted across the receptor structure (**Fig. 3B**).

**Fig. 3.**
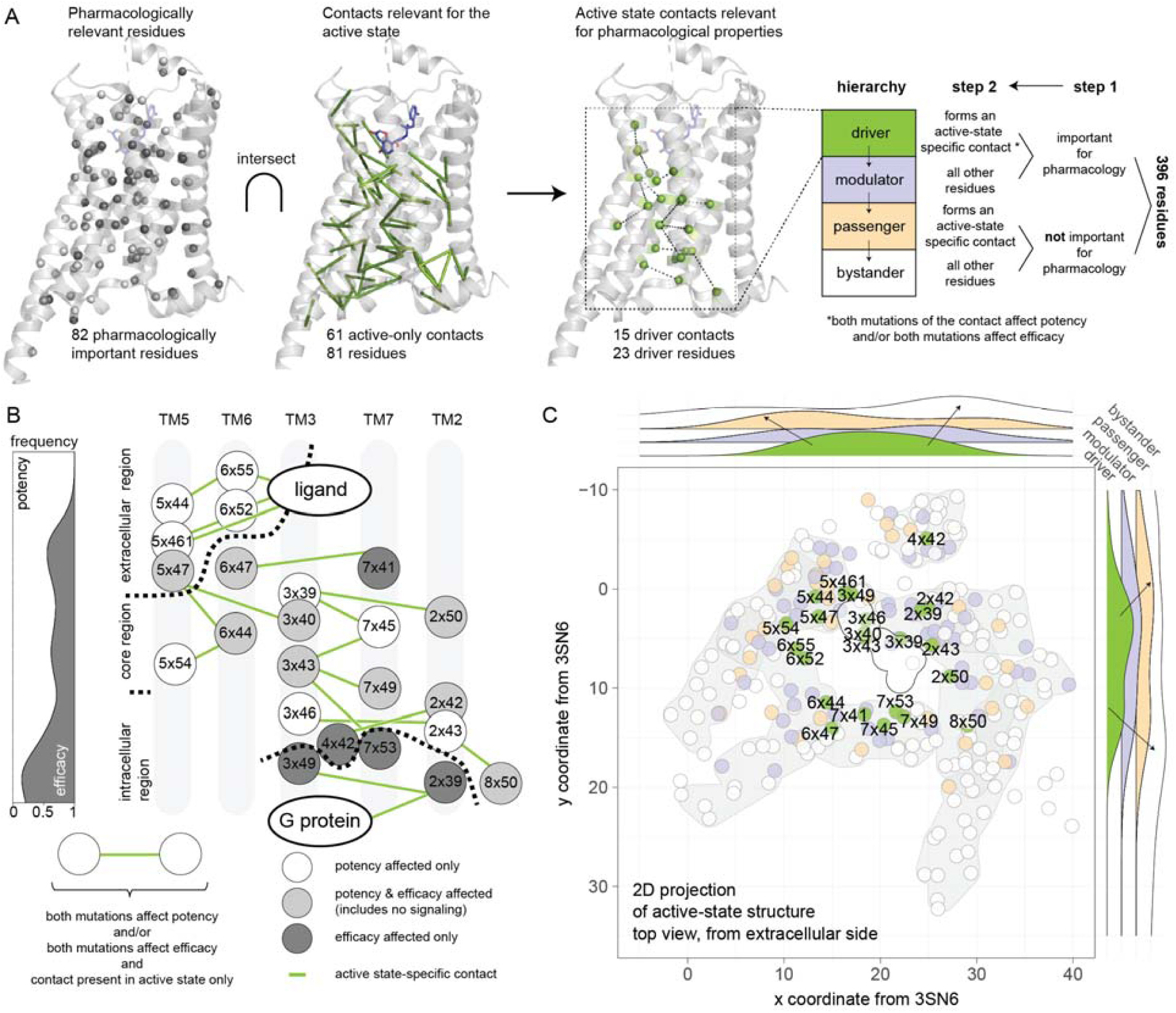
The network for efficacy and potency as determined by structure and pharmacology. **(A)** Integration of pharmacologically important residues and the residues contributing to newly established contacts in the active structure yields different classes of residues (see **main text**). **(B)** Residues forming the allosteric networks for efficacy and potency is represented as a cartoon, colored by effect of the mutation (dark grey: efficacy, white: potency, light grey: both potency and efficacy). The graph (left panel) depicts the frequency of potency versus efficacy effects for residues in the network, projected along the extra-to intracellular axis. (**C**) Position of residues classified by their participation in active state-specific contacts and importance for pharmacology on a 2D top view projection of the active, G protein-bound structure of the β2AR (PDB 3SN6). Frequency of residues along the x- and y-axis of the receptor is shown on the outside. Arrows denote the direction of the peaks of the distribution for the residue classes. Driver residues tend to be in the center whereas bystander residues tend to be at the periphery.

Our approach revealed 23 driver residues connected by 15 non-covalent contacts, constituting allosteric networks associated with efficacy and potency (**Fig. 3A**); of these 23, 9 affected potency only, 5 affected efficacy only and 9 affected both properties. As a general pattern, driver residues tended to be buried inside the receptor core, whereas modulator and passenger residues were located increasingly further away from the center, in proximity with solvent or membrane-exposed positions (**Fig. 3C**). Potency-affecting driver residues were enriched in the extracellular side of the receptor involving TM5 and TM6, and the efficacy-affecting driver residues were enriched at the intracellular side of the receptor in TM7, TM2, and TM3 (**Fig. S6**). Driver residues that affected both efficacy and potency were enriched in the core of the receptor, connecting the potency-only and efficacy-only residues. Overall, the allosteric network involving driver residues started extracellularly in TM6 and formed several connections between TM5 and TM6, including with F282^6×44^ of the previously identified PIF motif involved in receptor activation (*30, 31*)(**Fig. 3B**) before reaching TM3 in the receptor core. TM3 further connected to TM7 and the most conserved residue in TM2, D79^2×50^ (*22*), which corresponds to the allosteric sodium binding motif (Fig. 3D/E)(*32*). Finally, the network reached residues in the intracellular region, including D130^3×49^ and Y326^7×53^ of the conserved DRY and NPxxY motifs in TM3 and TM7, respectively.

The allosteric networks associated with efficacy and potency included three residues directly located in the ligand-binding site (N293^6×55^, F290^6×52^, and S207^5×461^); the other 8 ligand-binding residues do not form active-state specific contacts. This observation highlights the different roles of the ligand-contacting residues. While all 11 ligand-binding residues affected efficacy and/or potency when mutated, the former set of residues (three drivers) appear to initiate allosteric changes associated with active-state specific contacts and the latter set of residues (eight modulators) stabilize ligand-receptor interactions (**Fig. 3C, Fig. S6**). Consistent with this finding, the former set of residues that are involved in active state specific contacts (driver residues N293^6×55^, F290^6×52^, and S207^5×461^) make more contacts with other receptor residues than with epinephrine; In contrast, residues that are not involved in active state specific contacts (e.g., modulator residues D113^3×32^ and N312^7×38^) make substantial contacts with epinephrine and fewer contacts with other receptor residues (**Fig. S4**). Focusing on the G protein-binding residues, of the 10 that affected efficacy or potency, only one (driver residue T68^2×39^) is part of the allosteric network associated with efficacy and potency. These results suggest that the G protein-contacting residues that affect efficacy and potency can be classified into those disrupting the interaction with the G protein and the one that participates in the allosteric network (**Fig. 3B**). Residues of the allosteric networks in the core showed a tendency to affect either potency only or both potency and efficacy (**Fig. 3B**) and were enriched in TM3. These detailed structural and quantitative observations provide insight into the mechanism by which efficacy and potency emerges through structural and conformational rearrangements in β2AR.

While 23 of the 82 residues that affected efficacy and/or potency participate in active state-specific contacts, the remaining 59 (modulator) residues do not and hence are not part of the allosteric network associated with efficacy and potency. Some of these residues contact each other irrespective of the conformational state, forming 18 contacts between 27 residues. We categorized each of the 59 modulators according to their likely function (**Fig. S7**): 19 residues either contact the ligand directly or another residue that contacts the ligand or they lie in the putative ligand entry pathway; 16 are neighbors of driver residues either in sequence (i.e., within four residues) or in space (i.e., form a contact); 10 contact the G protein; and 8 may mediate lipid contacts in different places through aromatic or charged interactions. The roles for the remaining 6 residues were less clear and their determination may require further analyses on receptor stability and dynamics. Taken together, these results highlight how residues around allosteric networks may modulate ligand efficacy and potency in β2AR. The identification of modulator residues that lie on the receptor surface and interface with the allosteric networks, suggests possible structural sites for altering receptor signaling that can be targeted by allosteric ligands (*33*).

### Drivers, modulators, and passengers are under differential selection pressure

To assess the importance of the residues that differently affect pharmacology and/or active-state conformation, we performed an Evolutionary Trace (ET) analysis. ET is a phylogenetic method that identifies functionally important positions in protein families. In this analysis, low ET scores indicate positions that are highly conserved across homologs. Depending on how the alignments are constructed, residues with low ET scores can also represent positions that are conserved in a sub-family specific manner. We used the alignment of all adrenergic receptors and compared them with aminergic GPCRs to obtain the ET scores for each residue in the alignment (*34-36*). We then analyzed the distribution of ET scores for each of the residue categories defined above (driver, modulator, passenger, bystander). We find that driver residues are the most conserved, followed, in that order, by modulator, passenger, and bystander residues (**Fig. 4A**). These findings reveal that categorizing residues based on their pharmacological effects upon mutation and their role in mediating active-state specific contacts are associated with positions under purifying selection within the adrenergic and aminergic family of GPCRs. This association reflects the functional importance of these residue categories.

**Fig. 4.**
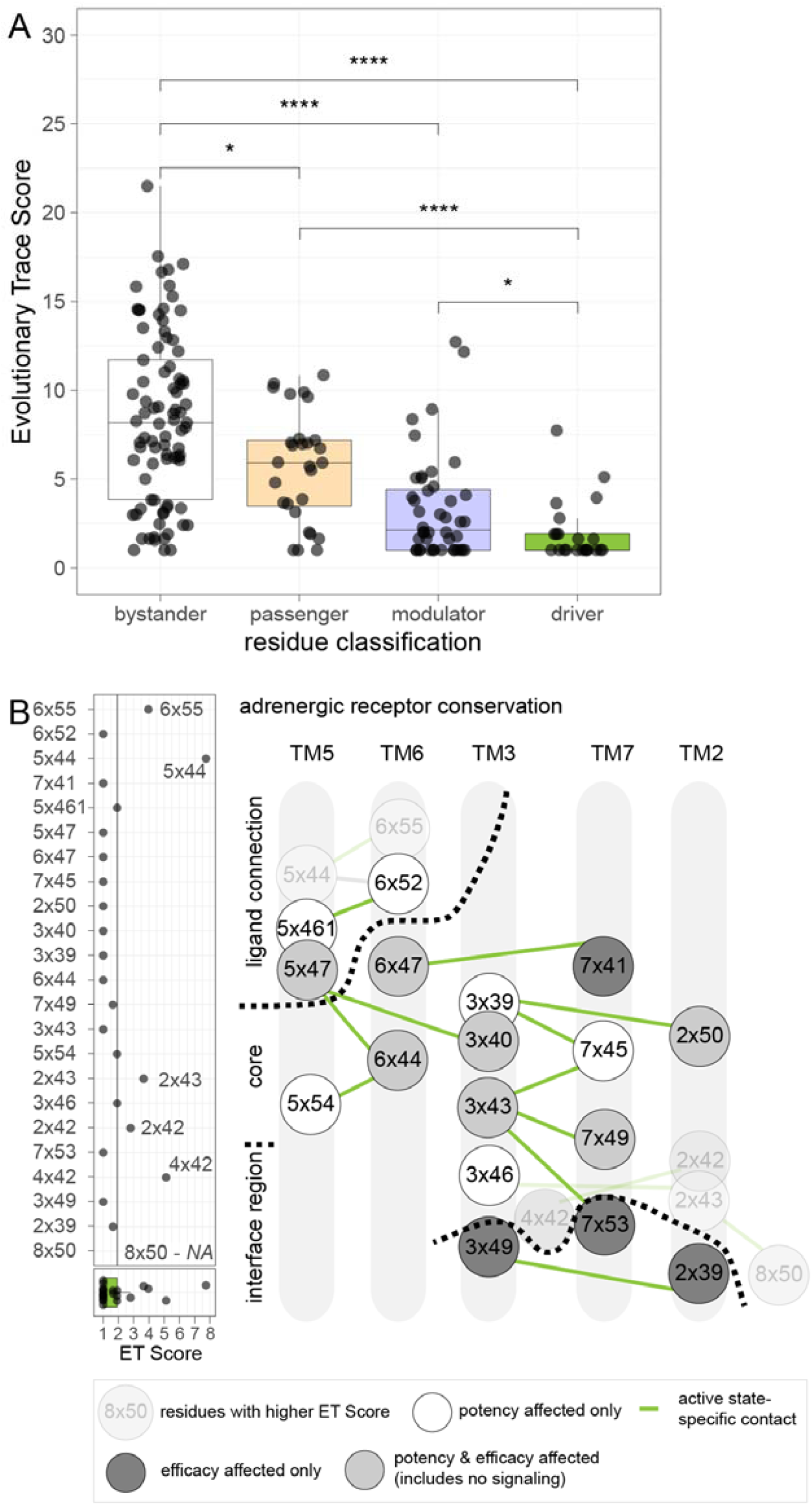
Residue conservation in adrenergic and aminergic receptors. (**A**) Adrenergic receptor ET score for residues for the different residue classes. Groups were compared using a Wilcoxon test (* p ≤ 0.05, ** p ≤ 0.01, *** p ≤ 0.001, **** p ≤ 0.0001) (**B**) Adrenergic ET scores for all network residues (left panel); highest ET scores (lowest conservation within adrenergic receptors) are labeled and made transparent on the cartoon representation of the allosteric network (right panel).

We then mapped the ET scores onto the allosteric networks associated with efficacy and potency to understand their distribution and position on the structure relative to functional regions. Driver residues with the lowest ET scores were at the center of the network in the receptor core, including all the network residues in TM 3 and TM7 (**Fig. 4B**). To contrast this finding, we determined which network positions showed the lowest conservation (i.e., higher variability), by extracting the network residues with the highest ET scores (> 3rd quartile, higher than 75% of the values). These more evolvable driver residues were either close to the ligand-binding site, or in the intracellular region (**Fig. 4B**), positions that may evolve or have evolved to recognize new ligands or G proteins.

Taken together, our findings highlight the differential selection pressure on pharmacologically important residues (driver and modulator residues), residues establishing active-state only contacts (passenger residues) and those that do not affect pharmacology or the active-state conformation (bystander residues) (**Fig. 4B**, for aminergic/family A receptors: **Fig. S8**).

## Discussion

Ligand efficacy and potency are the key pharmacological properties used to describe ligand-mediated signaling behavior. Yet, we do not fully understand how such pharmacological properties emerge from the structural rearrangements in a receptor, and how each amino acid influences ligand-mediated signaling. In this work, we have designed an integrative approach, combining pharmacological, structural, and evolutionary analyses, to investigate how GPCR sequence and structure encode a receptor’s ability to respond to an agonist.

To do so, we studied the β2AR, a well-studied member of the adrenergic receptor family and the larger family of aminergic receptors (*37*). There are over 30 high-resolution structures in the inverse agonist/antagonist-bound, agonist-bound and agonist/G protein-bound states (*27, 28, 38-40*), which have uncovered the large-scale and concerted conformational changes during the transition between the inactive and active states. Despite those detailed studies, how agonist binding and the resulting changes in the receptors’ intramolecular contacts relate to key pharmacological properties like efficacy and potency remains unclear. Similarly, mutating selected residues and subsequent determination of signaling responses have provided valuable insights into the relevance of specific residues (*20, 41*), but we still do not fully understand the structural basis for the observed signaling differences.

To address these challenges, we have comprehensively analyzed how β2AR residues involved in conformational transitions are associated with receptor activation and G protein signaling in response to epinephrine, an endogenous agonist for this receptor. By developing a computational approach to integrate systematic mutagenesis and pharmacological analyses with data on the conformational changes that occur during receptor activation, we have been able to reveal the allosteric networks associated with efficacy and potency. To our knowledge, such detailed analyses have not been performed for any GPCR and allow us to provide a more nuanced view of receptor activation.

We identify a core set of 23 driver residues and 59 modulator residues that influence epinephrine efficacy and potency in β2AR. The driver residues are interconnected to form a network of active state-specific contacts that spans from the extracellular to the intracellular regions of β2AR. Most importantly, by explicitly considering efficacy and potency readouts in the analysis, we provide new insights into β2AR signaling, revealing a spatial segregation of residues affecting potency and efficacy in the active-state specific network and insights into the likely function of the modulator residues. Furthermore, analysis of the active-state specific residues contacts in the context of the pharmacological data allows us to identify 58 passenger residues. Mapping these residue classifications onto the structure reveals that driver residues are buried in the receptor core, with modulator and passenger residues positioned around them. Consistent with these observations, an evolutionary trace analysis revealed that driver residues are most conserved, followed by the modulators, passengers, and bystander residues.

The residues, their roles and the network defined here are specific for the particular ligand, receptor, and effector system (i.e., epinephrine, β2AR and Gs, respectively). For the same receptor, we expect that the residues and the networks will slightly deviate according to the specific ligand and the signaling partner, including the specific G protein sub-type. Performing similar analyses with arrestin instead of G protein, should yield new insights into the difference in activation mechanism of G proteins and arrestins. Future analyses of inactive and G protein-or arrestin-bound structure pairs in the context of the relevant pharmacological data will shed light onto the similarities and differences in networks and the residues required for eliciting the complete pharmacological response. We envision that the framework developed here will allow investigation of such questions in the future, leveraged by the development of high-throughput approaches to pharmacologically characterize mutations and the increasing availability of new structures.

In conclusion, our work reveals a GPCR activation network and the contributions of specific residues towards ligand potency and efficacy. Apart from revealing general principles, the approach and the findings presented here can help understand how disrupting parts of the network could alter receptor output. In this manner, our work has implications for the way we select, discover, and rationalize the effects of new orthosteric and allosteric GPCR ligands, depending on how they engage with parts of the network. It could also help understand the role of lipid-interacting residues and suggest how changes in particular receptor residues in the form of missense natural variants or disease-related mutations could influence signaling responses to endogenous agonists or GPCR drug treatments.

## Acknowledgments

We would like to thank Ines Chen and John Janetzko for reading the manuscript and providing comments. We thank members of Kobilka and Babu groups for constructive feedback on this work.

## Funding

Marie Skłodowska-Curie Individual Fellowships from the European Union’s Horizon 2020 research and innovation programme under grant agreements No 844622 and 832620 (FMH, MMS).

American Lebanese Syrian Associated Charities (ALSAC; MMB)

American Heart Association Grant # 19POST34380839 (FMH).

Federation of European Biochemical Societies Long-Term Fellowship (MMS).

National Institutes of Health grant R01NS028471 (BKK)

Canadian Institute of Health Research grant FDN#148431 (MB)

UK Medical Research Council grant number MC_U105185859 (MMB)

MB holds a Canada research Chair in Signal Transduction and Molecular Pharmacology

MMS is a Wolfson College Junior Research Fellow

BKK is a Chan Zuckerberg Biohub Investigator.

## Author contributions

Conceptualization: FMH, BKK, MB, MMB

Methodology: FMH, MB, MMB

Formal analysis: FMH

Investigation: FMH, MMS, MS

Project administration: FMH

Visualization: FMH

Software: FMH

Funding acquisition: FMH, MMS, BKK, MMB, MB

Supervision: BKK, MMB, MB

Writing – original draft: FMH, MMS, MS

Writing – review & editing: FMH, MMS, MS, BKK, MMB, MB

## Supplementary Information

### Materials and Methods

#### DNA constructs and mutagenesis

The human β2-adrenergic receptor (β2AR) was codon-optimized and cloned into pcDNA3.1 together with an N-terminal signal sequence, Twin-Strep-tag, and SNAP-tag. BRET-based biosensor constructs for Gs, Gβ1, and Gγ1 were in pcDNA3.1. The Gs biosensor consisted of RlucII-Gαs, where Renilla luciferase (RlucII) was inserted after amino acid 67, unmodified human Gβ1, and GFP10-Gγ1.

β2AR mutants for positions 2-412 were generated as described (*42*). Amino acids other than alanine were replaced by alanine, glycines replaced native alanines. Briefly, primers were designed using the custom-made software AAScan (*43*) (available here: https://github.com/dmitryveprintsev/AAScan) and ordered from Integrated DNA Technologies (IDT) as 500 pmol DNA oligos in 96-well plates. Forward and reverse mutagenesis primers were used in separate PCR reactions together with one primer each, which annealed in the ampicillin resistance gene (CTCTTACTGTCATGCCATCCGTAAGATGC and GCATCTTACGGATGGCATGACAGTAAGAG). PCRs were run as touchdown PCRs in 20 μl final volume. Resulting half-vector fragments were combined, digested with 0.5 μl DpnI (NEB) for 1 h, cleaned up using ZR-96 DNA Clean & Concentrator-5 (Zymo Research), assembled by Gibson assembly using HiFi DNA assembly Master Mix (NEB) at 45°C for 1 h (*44*) and transformed using Mix & Go competent cells (Zymo Research). The resulting colonies were cultured in 5 ml LB medium in 24-well plates. The DNA was purified using Qiaprep 96 Plus kits (Qiagen) and sent for sequencing (Bio Basic Inc. or in-house). Sequences were analyzed using custom software MutantChecker (available here: https://github.com/dmitryveprintsev/AAScan). Mutants that were not successfully generated as described were cloned using an alternative method: pcDNA3.1 was digested with NheI and XhoI (NEB), the β2AR coding sequence was amplified in two PCRs, each using one of the mutagenesis primers and either T7long or BGH primer (CGAAATTAATACGACTCACTATAGGGAGACCCAAGCTGG and TAGAAGGCACAGTCGAGG) which anneal upstream and downstream of the coding sequence, respectively. The two fragments were combined, digested using DpnI and cleaned up as described above. The clean fragments were assembled with digested, cleaned pcDNA3.1 using HiFi DNA assembly Master Mix (NEB).

#### BRET-based signaling assays

HEK-293 SL cells (a gift from Stéphane Laporte) were cultured in DMEM supplemented with 4.5 g/l glucose, L-glutamine, and 10% newborn calf serum (NCS, Wisent BioProducts, Canada) and penicillin-streptomycin (PS, Wisent BioProducts, Canada) at 37°C with 5% CO_2_. Cells were transiently transfected using linear 25 kDa polyethyleneimine (PEI, Polysciences Inc., Canada, No. 23966) in a 3:1 ratio with DNA. Cells were seeded at 20’000 cells per well into Cellstar® PS 96-well cell culture plates (Greiner Bio-One, Germany) and incubated for two days at 37°C, 5% CO_2_. Before measurement, the cells were washed with PBS followed by the addition of Tyrode’s buffer (137 mM NaCl, 0.9 mM KCl, 1 mM MgCl_2_, 11.9 mM NaHCO_3_, 3.6 mM NaH_2_PO_4_, 25 mM Hepes, 5.5 mM glucose,1 mM CaCl_2_, pH7.4) and incubated for at least 30 min at 37°C. Plates were treated with ligand for 10 min before measurement and with coelenterazine 400a (Nanolight Technology) at 5 μM final 5 min prior to measurement. Ligand concentrations used were: 31.6 nM (10^−8.5^ M) to 3.16 mM (10^−3.5^M) in half-log steps plus a buffer control.

Coelenterazine 400 a was prepared with 1% Pluronic F-127 in Tyrode’s buffer to increase solubility. Epinephrine was prepared in 0.01 M HCl to increase stability, the low pH was buffered by Tyrode’s buffer upon addition of ligand to the cells. The BRET signal was read in a Synergy Neo (Biotek) equipped with dual photomultiplier tubes (PMTs, emission: 410 nm and 515 nm, gain 150 for each PMT and 1.2 s integration time).

All signaling experiments were done in biological triplicates, wild-type and mock transfection controls were repeated three times per measurement day.

#### Cell-surface ELISA

HEK-293 SL cells were transfected using PEI as above and seeded into poly-L-lysine coated 96-well plates (Greiner BioOne, Germany) and incubated for 2 days at 37°C 5% CO_2_. Each well was washed with 200 μl PBS, the cells were then treated with 50 μl 3% paraformaldehyde per well for 10 min. Each well was washed as follows: Three washes with wash buffer (PBS + 0.5% BSA); in the last step, the cells were incubated with wash buffer for 10 min. Primary rabbit anti-SNAP antibody was added at 0.25 µg/ml in 50 μl and incubated for 1 h at RT, followed by three wash steps as described above. The cells were incubated with secondary anti-rabbit HRP antibody (GE Healthcare) diluted 1:1000 for 1 h. Again, the cells were washed three times with wash buffer and then three times with PBS. Per well, 100 µl SigmaFast solution was added, the plates were incubated at RT in the dark. Reactions were stopped by the addition of 25 μl 3M HCl. 100 μl of the resulting solution was transferred to a new 96-well plate (transparent), and the optical density was read at 492 nm using a Tecan GENios Plus microplate reader. Cell-surface ELISAs were carried out in three biological replicates with internal quadruplicates. Each plate contained wild-type and mock-transfected cells as controls.

#### Data analysis

##### Processing of signaling data

The BRET data were processed using BRET2DTF and DataFitter custom software (Dmitry Veprintsev; https://github.com/dbv123w/DataFitter). All concentration-response curves were fitted to a Hill equation using a Hill slope of 1 to determine the agonist concentration at the half-maximal signal response (logEC50) as well as the pre-transition and post-transition baselines. All concentration-response curves were visually inspected and manually curated.

The results from data fitting were read into R (RStudio 1.3.959 and R 4.0.0) and processed using custom scripts. The following packages were used: tidyverse (especially dplyr, ggplot2, purrr, tibble, tidyr, forcats, stringr), plotly, MASS, reshape, reshape2, ggrepel, patchwork, ggpubr, bio3d (*45*), openxlsx.

The data were read in, reformatted, averaged, annotated (addition of GPCRdb number, expression level, amino acid number, mutation) filtered based on the results of manual curation, normalized, and visualized in R. We checked the data set for outliers, day-to-day variation and trends, the effect of expression level on signaling response and baseline signal variation. Measured mutant data were normalized to wild type using the most recently measured wild type data set to correct for day-to-day variation and variation during the day. Cut-offs were applied to the normalized, corrected, and curated data, resulting in discretization. These cut-offs were chosen so that none of the signaling measurements from a total of 95 wild-type concentration-response curves were classified as not wild-type like. For the normalized amplitude, the cut-off was 0.74 (where wild-type is centered around 1), and for the normalized logEC50, the cut-off was 0.87 (where wild-type is centered around 0); this corresponds to a 7.4-fold change. Graphs were generated in ggplot2 (part of the tidyverse) and post-processed in Adobe Illustrator CS6.

##### Residue-residue contacts

Residue-residue contacts were obtained from Protein Contacts Atlas (https://www.mrc-lmb.cam.ac.uk/rajini/index.html) (*29*). Two residues are listed as a contact if the distance between the two atoms minus their van der Waals radii equals 0.5 Å or less. Text files downloaded from Protein Contacts Atlas containing residue-residue contacts were read into R and cleaned. We then fortified the tables with additional information (secondary structure elements (SSE) and GPCRdb numbers for each amino acid), filtered the contact tables to exclude main chain-main chain contacts, and include contacts between two different SSEs only. The contacts were filtered to include only those residues resolved in both crystal structures used for the comparison, 2RH1 and 3SN6. In the inactive and in the active G protein-bound structures of the β2AR, 282 and 285 residues form 1275 and 1233 residue-residue contacts, respectively. To focus on residue contacts that contribute to large domain motions instead of local rearrangements during activation, we excluded contacts formed exclusively between backbone atoms and contacts within the same secondary structure element (i.e., contacts formed within a single helix). This reduced the number of residue-residue contacts to 117 unique contacts in the inactive state, 61 unique contacts in the active, G protein-bound state, and 191 contacts that are present in both the active and inactive states.

##### Definition of the ligand and G protein binding sites

Definition of the ligand-binding site and the G protein binding site were based on the structures of β2AR with epinephrine and an engineered nanobody (PDB 4LDO) (*21*), in complex with heterotrimeric G protein and Gs peptide (3SN6 and 6E67) (*27*) (*46*). All residues within 4Å of epinephrine were classified as part of the ligand-binding site, the residues were verified using LigPlot+ v.2.2 and literature data (*22, 47*) (*48*) (*49*) (*50*). All residues within 4 Å of G protein or Gs peptide were classified as residues in the G protein binding site.

##### Residue classification

We defined driver and modulator residues as those residues that strongly affect Gs signaling upon mutation. Additionally, driver residues also form an active-state specific contact to another residue. Both residues that mediate such a contact must share an effect on Gs signaling, that is, either both residues negatively affect potency, or both affect efficacy. Residues not forming such contacts but affecting pharmacology were classified as modulators. Passenger and bystander residues were defined as those residues that do not affect Gs signaling upon mutation. Additionally, passenger residues must form at least one active-state specific contact. Residues not forming such contacts and not affecting Gs signaling upon mutation were classified as bystanders. Hierarchically, we defined drivers as the structurally and pharmacologically most important residues, followed by the pharmacologically important modulators, the passenger residues involved in conformational change but not pharmacologically important and finally the bystanders that are not involved in conformational changes and not important pharmacologically.

**Fig. S1.**
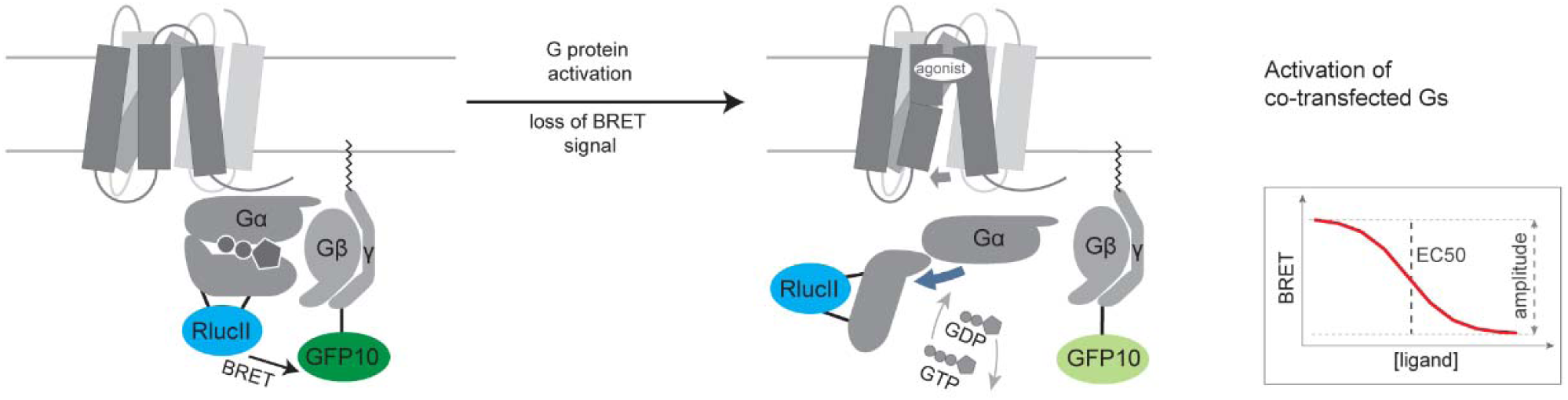
BRET-based biosensor for Gs activation. To measure Gs activation in HEK cells, we used a BRET-based biosensor for Gs activation. The Gα subunit was fused with *Renilla reniformis* luciferase (RlucII, donor) and the Gγ subunit was labeled with GFP10 (acceptor). Upon G protein activation, the Gα subunit changes its conformation and dissociates from Gβγ, leading to a decrease in BRET signal.

**Fig. S2.**
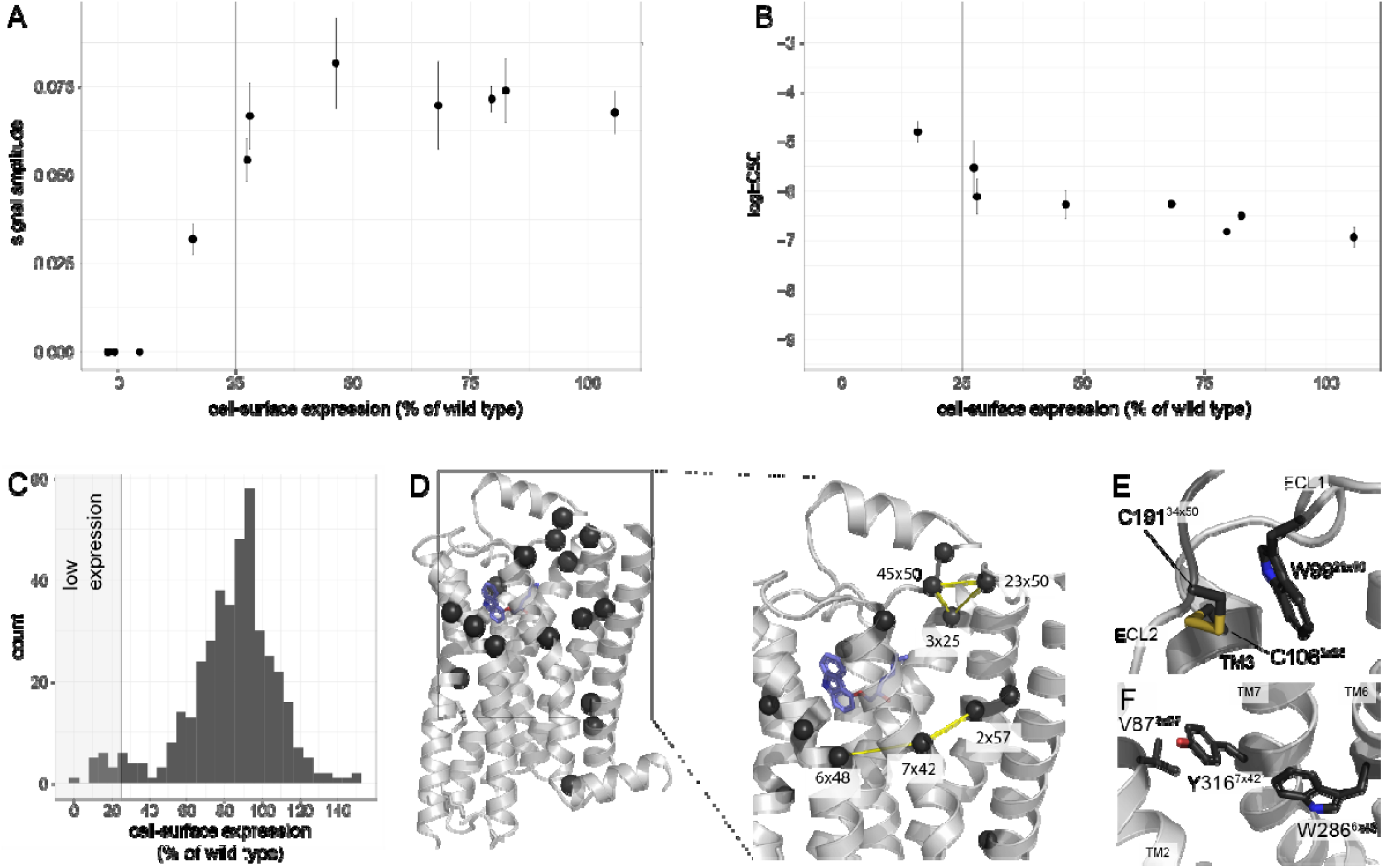
Residues important for cell-surface expression. (**A**) Variation of the signal amplitude with cell-surface expression (**B**) Variation of the logEC50 with cell-surface expression (**C**) Cell-surface expression level of 412 single point mutations shown as a histogram. Low expression was defined as <25% of wild-type expression level. (**D**) Position of residues whose mutation led to a strong decrease in cell-surface expression is shown as black spheres. These residues formed two networks: one involving the extracellular disulfide bridge (C106^3×25^ and C191^34×50^) and a tryptophane (W99^23×50^), the other connected helices 6, 7, and 2 (V87^2×57^, W286^6×48^, and Y316^7×42^). (**E**) and (**F**) stick representation of the two networks.

**Fig. S3.**
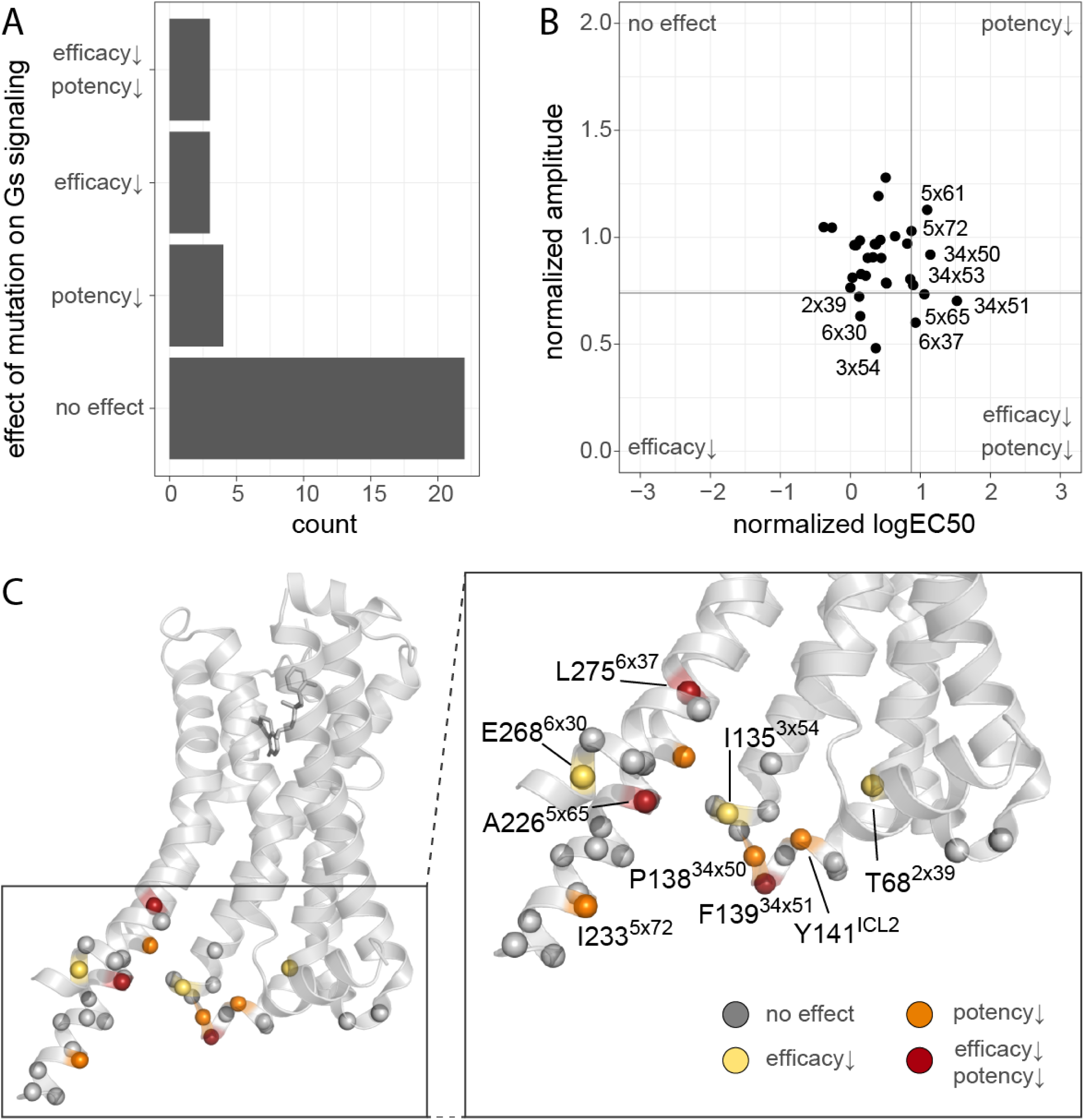
Effect of mutating residues in the G protein binding site. (**A**) Residue counts by effect on signaling. (**B**) normalized amplitude and logEC50 for all residues within 4 Å of Gs in the β2AR-Gs structure (PDB: 3SN6) and/or G peptide in the fusion structure with β2AR (PDB: 6E67). (**C**) Residues in the Gs binding site shown on the β2AR-Gs structure (PDB: 3SN6) are shown as spheres, colored by effect: no effect (gray), reduced potency only (orange), reduced efficacy only (yellow) and both reduced efficacy and potency (red).

**Fig. S4.**
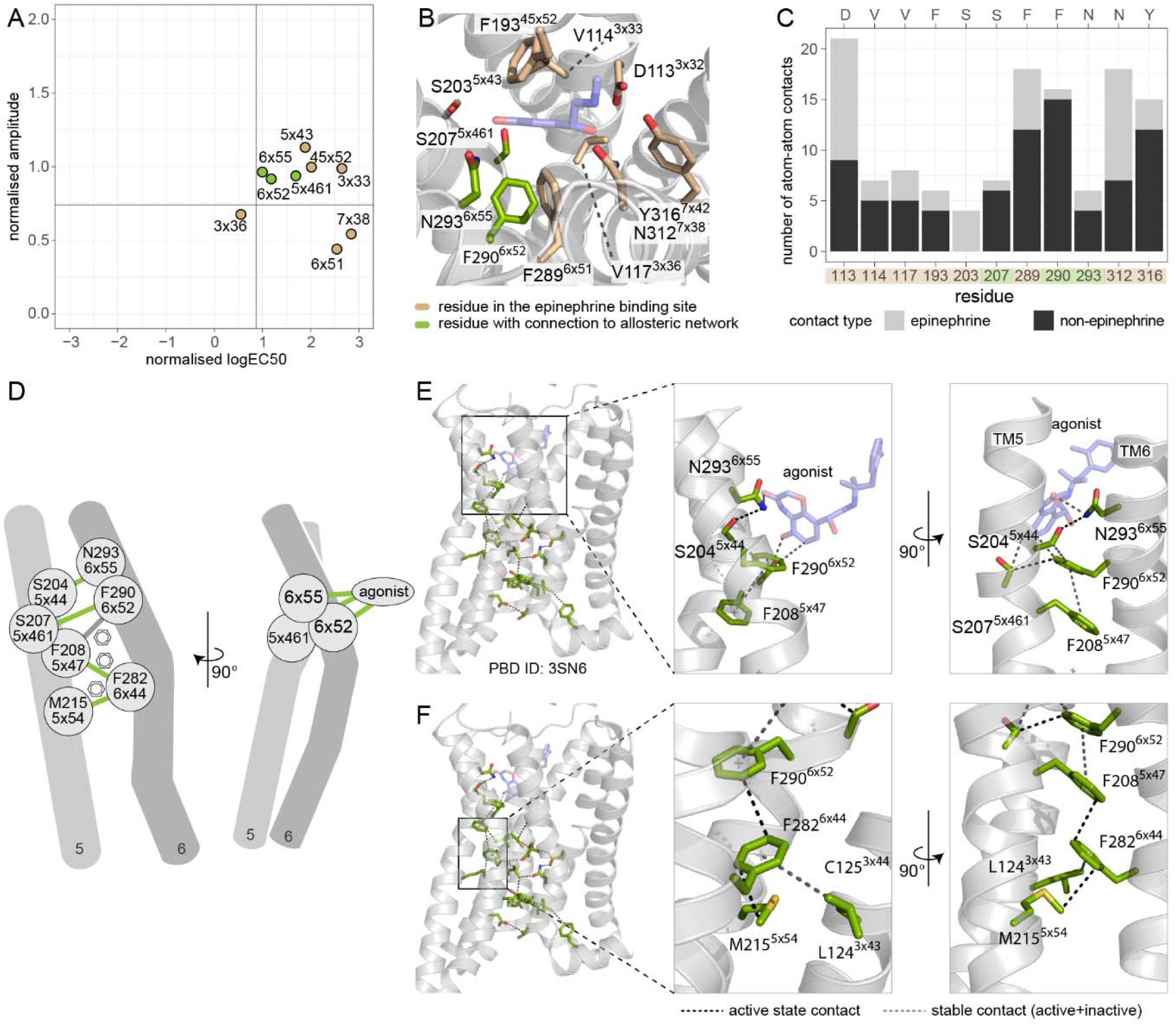
Residues 6×55, 6×52, and 5×461 connect the ligand-binding site to the signal transduction network. (**A**) Effect of mutation for residues in the epinephrine binding site. (**B**) Structural representation of the epinephrine binding site (PDB 4LDO). (**C**) Number of atom-atom contacts for binding site residues, colored by contact type (**D**) Residues connecting ligand and receptor core with active-state specific contacts (**E** and **F**) detailed view of the extracellular side of the network.

**Fig. S5.**
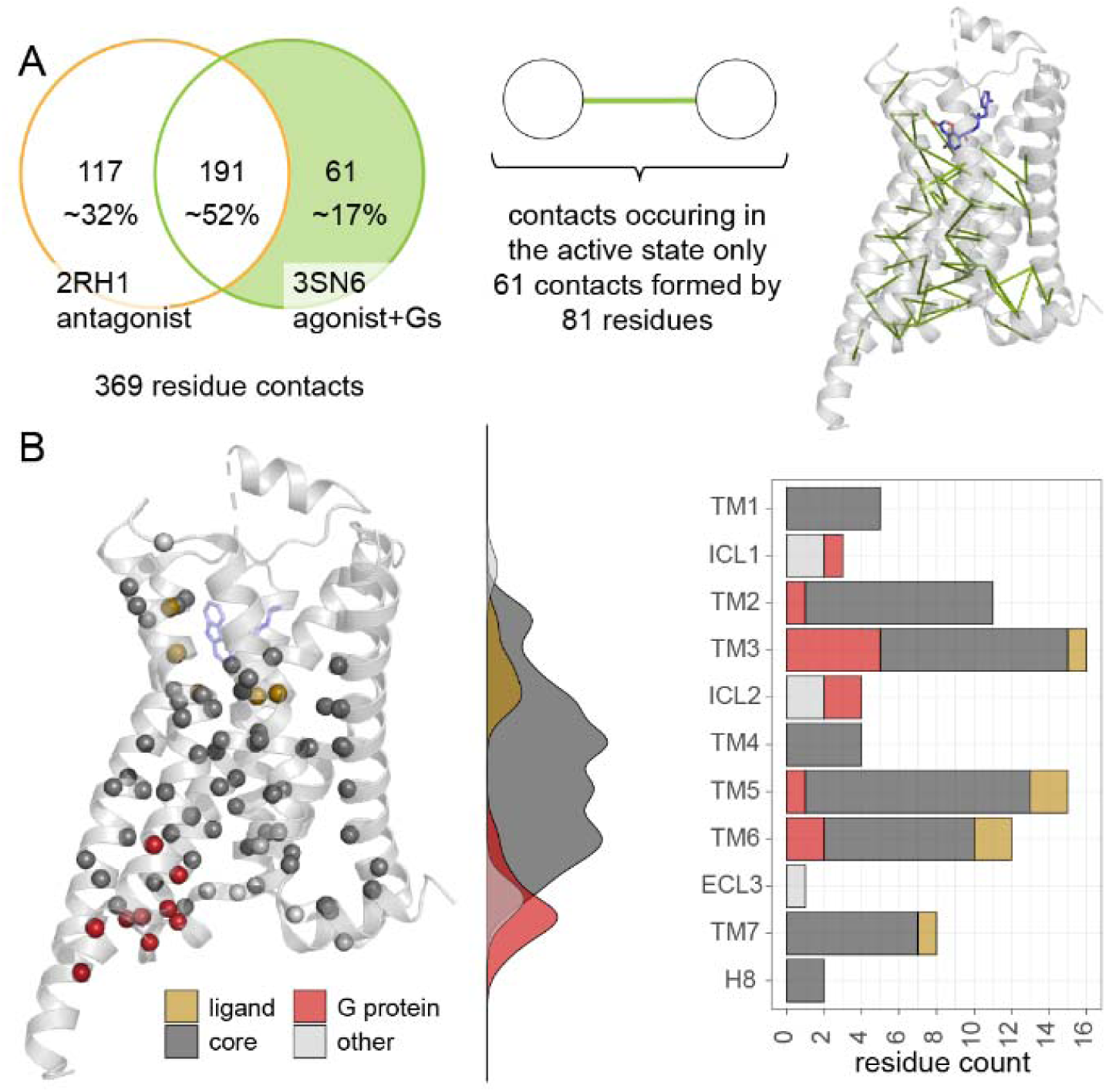
Residues that form active-state specific contacts. (**A**) Contacts specific to the active, G protein-bound state based on the structures of the β2AR bound to carazolol (inactive, 2RH1) and BI-167107 (G protein complex, 3SN6) and the position of the contacts in the 3SN6 structure. (**B**) Distribution of residues forming active-state specific contacts, colored by functional site (ligand-binding site (yellow), G protein binding site (red), core (dark grey), other (light grey)).

**Fig. S6.**
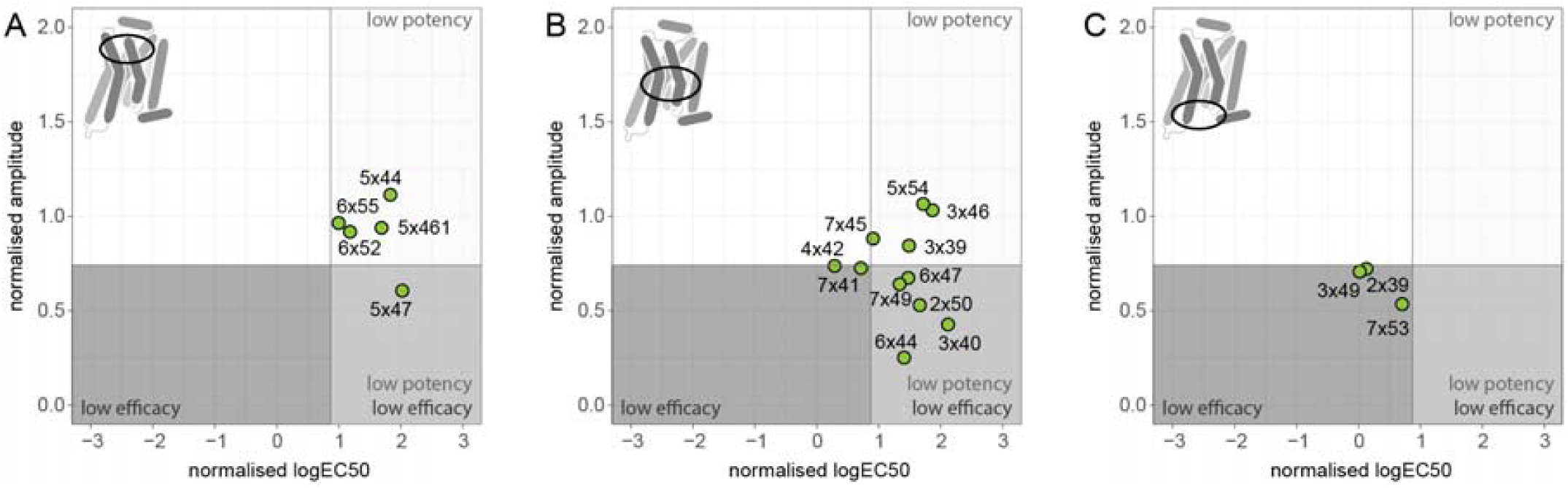
Effect of mutating residues in the allosteric network on Gs activation. (**A**) Effect of mutating residues in the extracellular region (**B**) Network residues in the receptor core (**C**) Network residues in the intracellular region (defined as any residue within 10 Å of G protein).

**Fig. S7.**
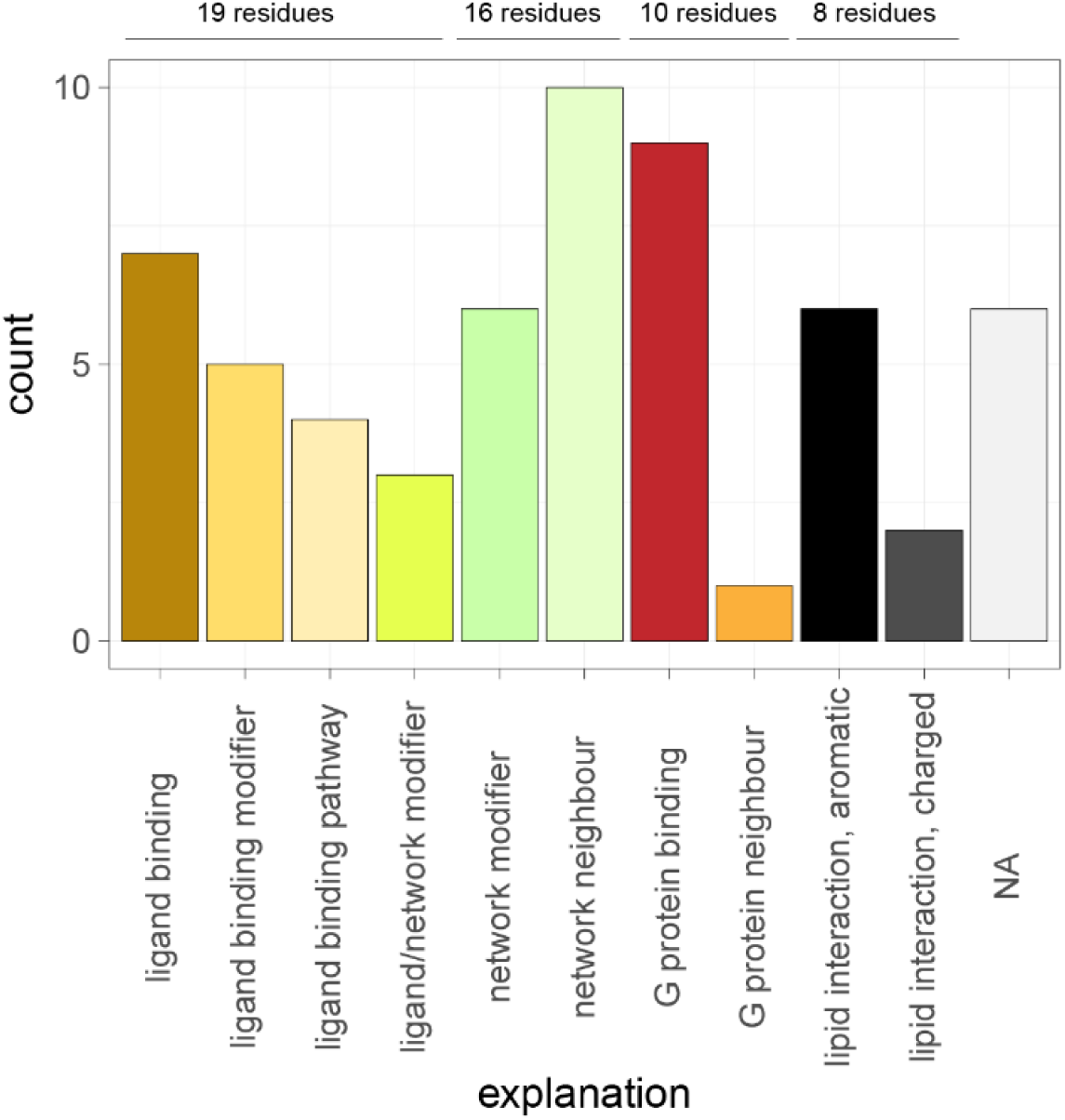
Potential role of pharmacologically important modulator residues. Ligand binding residues are residues within 4 Å of epinephrine (PDB 4LDO); driver residues were excluded in the graph. Residues likely to be in the ligand’;s proximity during the ligand-binding process were termed ligand-binding pathway. Modifiers were defined as residues within proximity of residues in the network, the binding site, or both. Neighbors are residues within 4 residues of the indicated functional site in the same secondary structure element. Functional assignments were done manually.

**Fig. S8.**
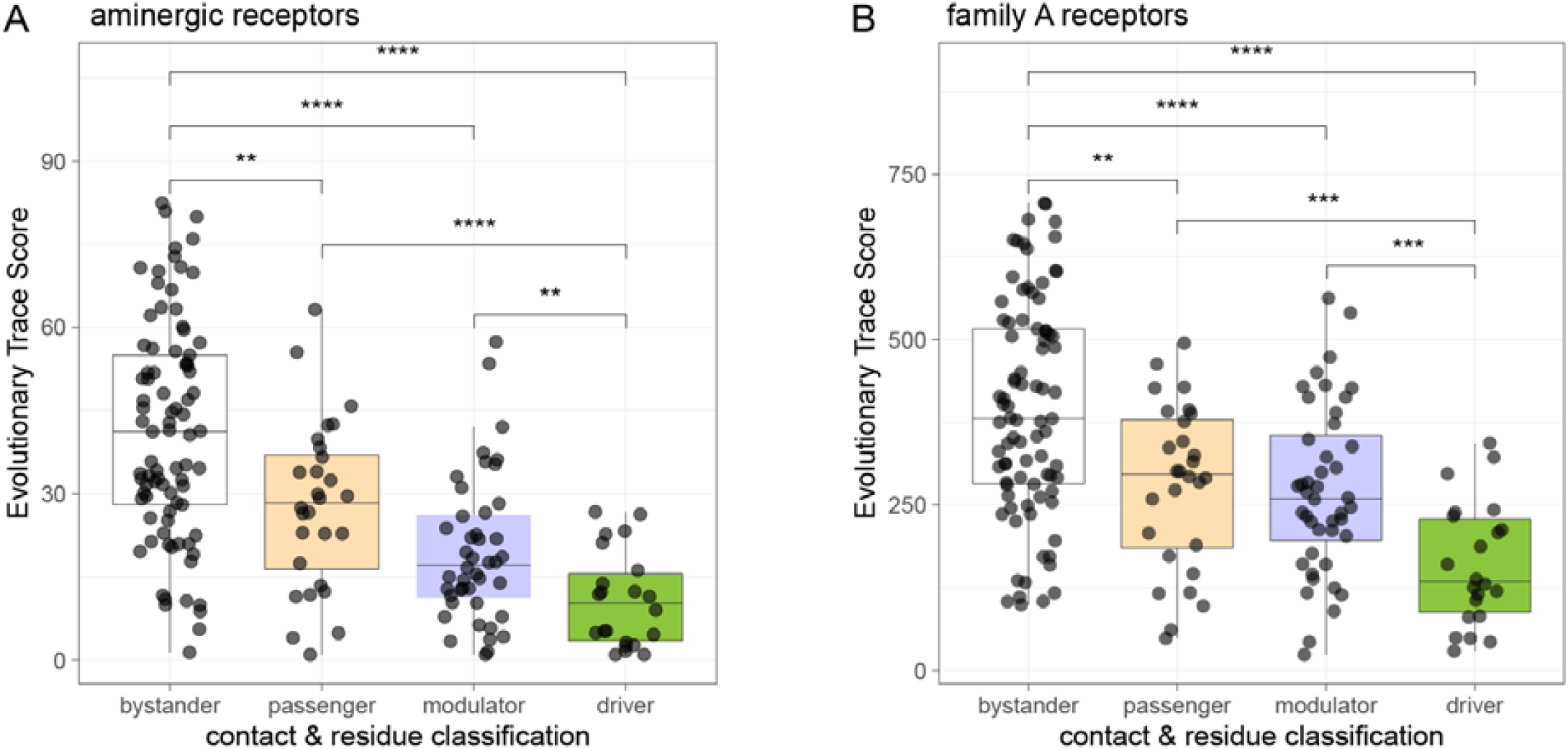
Evolutionary trace analysis of amino acid conservation of network residues. (**A**) Evolutionary Trace analysis for aminergic receptors. Groups were compared using a Wilcoxon test (* p ≤ 0.05, ** p ≤ 0.01, *** p ≤ 0.001, **** p ≤ 0.0001) (**B**) Evolutionary Trace analysis for 5105 family A GPCR sequences (not limited to human sequences).

